# Reversal of drug resistance by disruption of a Gain-of-Function mutant p53 and transcriptional co-activator PC4 interaction

**DOI:** 10.1101/2023.02.17.528954

**Authors:** Priya Mondal, Kumar Singha Roy, Tapas K. Kundu, Susanta Roychoudhury, Siddhartha Roy

**Affiliations:** Department of Biophysics, Bose Institute, P-1/12 C.I.T. Scheme VII M, Kolkata 700054, India; Cancer Biology and Inflammatory Disorder Division, CSIR-Indian Institute of Chemical Biology, 4 Raja S.C. Mullick Road, Kolkata 700032, India; Transcription and Disease Laboratory, Molecular Biology and Genetics Unit, Jawaharlal Nehru Centre for Advanced Scientific Research, Bangalore, 560064, Karnataka, India

**Keywords:** Drug resistance, p53, Gain-of-function, Peptide, Protein-protein interaction

## Abstract

The positive coactivator 4 or PC4 is a chromatin-associated protein whose role in gene regulation by wild-type p53 is now well-known. During tumorigenesis, p53 is often mutated resulting in its loss of function. A sub-class of these mutants gain new pro-proliferation properties which occur largely due to the upregulation of many pro-proliferation genes. Little is known about the roles of PC4 in tumor cells bearing mutant p53 genes. In this article, we show that PC4 associates with one of the tumor-associated gain-of-function p53 mutants, R273H. This association drives its recruitment to two promoters, UBE2C, and MDR1, known to be responsible for imparting aggressive growth and resistance to many drugs. A previously reported peptide that disrupts PC4-wild-type p53 interaction also disrupts the PC4-R273Hp53 protein-protein interaction. The introduction of this peptide to tumor cells bearing the R273HTP53 gene resulted in a lowering of MDR1 expression and abrogation of drug resistance. Interestingly, cells bearing another gain-of-function mutant R248W do not show the same type of response, suggesting that the action of PC4 on mutant p53s may differ for different GOF mutants. The results presented here suggest that PC4-R273H interaction may be a promising target for reducing proliferation and tumor drug resistance.

---

The master transcription factor p53 puts up a major barrier against oncogenesis by regulating many well-known pathways that promote tumor-suppressive functions (*1*). In addition to the canonical tumor-suppressive pathways, p53 can oppose tumorigenesis by many other lesser-known mechanisms (*2*). Inactivation of p53 by mutations that cause loss of its transcription regulatory function is one of the major mechanisms that leads to a full-fledged malignant transformation. Within this set of loss-of-function mutations, a subset of mutations also gains additional pro-oncogenic functions by upregulating genes related to proliferation and drug resistance (*3*). This latter subset of mutations has been called gain-of-function mutations. Often these gained functions confer aggressive character to these cells.

How the gain-of-function mutant p53 protein upregulates the pro-oncogenic genes, is not known in detail. It appears now that some gain-of-function mutants piggyback on other pro-growth transcription factors and translocate to genes related to proliferation and drug resistance (*4*). Once recruited there, the gain-of-function mutants may upregulate the gene, perhaps through interaction with other protein factors.

Many aggressive tumors have gain-of-function p53 mutants which play a crucial role in their ability to evade therapeutic interventions. If these gain-of-function properties can be reversed by external agents, it may be possible to use many of the extant therapies to control or even reverse tumor growth. Several approaches have already been tried in the laboratory, with varying degrees of success (*5*). An often-used method is to try to convert the mutant p53 protein into the wild-type conformation, having properties of the wild-type protein (*6*). Here we take a different and novel approach. We target a previously unknown protein-protein interaction between a gain-of-function mutant of p53 and a chromatin-associated protein as well as a transcriptional coactivator, PC4, by disrupting their interaction using a small peptide, derived from the p53 sequence. Disruption of this interaction leads to the reversal of drug resistance making the cells susceptible to standard chemotherapeutic drugs.

## Materials and Methods

### Cell culture

Human lung cancer cell line H1299 was purchased from American Type Culture Collection (ATCC). PANC-1 was purchased from National Centre for Cell Science (Pune, India). H1299 was grown in Roswell Park Memorial Institute (RPMI) 1640 medium (Gibco). PANC-1 was grown in Dulbecco’s Modified Eagle’s Medium (DMEM) (Gibco). Mutant p53–expressing stable cell lines (H1299-R248W) and G418+ H1299-R273H were previously described (*7*). Both media was supplemented with 10% fetal bovine serum (Gibco). All cell lines were cultured at 37°C in a humidified 5% CO_2_ incubator. All cell lines were routinely screened and tested for mycoplasma contamination.

### Peptide Synthesis, labeling, and Purification

All peptide nomenclature and corresponding sequences are shown in Table S1. Peptide synthesis, labeling, purification and characterization have been previously described in detail (*8*). Expected and measured Molecular Masses are reported in Table S1 and the Mass spectra are reported in Figure S1.

### Semi-Quantitative PCR

All PCRs were performed by using Recombinant Taq polymerase (Thermo Scientific). 100 ng of the synthesized DNA was mixed with 1X PCR buffer, 1 µM reverse primer, 1 µM forward primer, 0.2 mM dNTPs, 2 mM MgCl_2_ and 1.25 units of Taq polymerase. The volume was adjusted to 25 µl with nuclease-free water. The reaction was set up in a thermocycler (GeneAmp 9700 PCR System, Applied Biosystems). The PCR products were resolved on 1.8% agarose (USB) gel and visualized by staining with ethidium bromide (USB). The primer sequences and conditions of PCR reactions are described in Table S2.

#### Quantitative real-time PCR

RNA isolation, cDNA synthesis and other methodologies have been described in Mondal *et al*. (*9*). Real-time PCR was performed in the 7500 Fast Real-Time PCR System (Applied Biosystems) with DyNAmo ColorFlash SYBR Green qPCR Kit (Thermo Scientific). 100 ng of the prepared cDNA was mixed with 1X SYBR Green PCR master mixes, 1 µg forward primer and 1 µg reverse primer. The volume of the reaction mixture was adjusted to 10 µl with nuclease-free water. The data were analyzed by 7500 Fast software (Version 2.3, Applied Biosystems). The comparative threshold cycle method (ΔΔCt) was used to quantify relative amounts of product transcripts with GAPDH as the endogenous reference control. The primer sequence and conditions of PCR reactions are described in Table S2.

#### Western blot

Cells were treated with different doses of peptides and anticancer drugs and incubated for time periods as required. Cells were lysed with radioimmunoprecipitation buffer (50 mM Tris, pH 7.5 containing 15 mM EDTA, 150 mM NaCl, and 1% NP-40) supplemented with protease inhibitor cocktail (Sigma Chemical Company) and PMSF (Sigma Chemical Company). Total protein concentration in each lysate was measured using the standard BCA (Thermo Scientific) method. 50 μg proteins were mixed with 20% v/v of 5X SDS lysis buffer and 10% β-mercaptoethanol and boiled at 100°C water bath for 5 minutes. The supernatant was collected by centrifugation at 13000 rpm for 10 min and loaded on PAGE (Poly-acrylamide Gel Electrophoresis).

Antibody-reactive bands were detected by enzyme-linked chemiluminescence. Chemiluminescence detection of bands was done with Super Signal West Pico Chemiluminescence substrate (Thermo Scientific). The HRP substrate consists of Luminol Reagent and Peroxide Solution. Working HRP substrate was prepared by combining equal volumes of Luminol Reagent (Substrate A) and Peroxide Solution (Substrate B) at room temperature. Approximately, 1 ml of working HRP substrate was required per 9cm × 6cm of the membrane area. The working HRP substrate solution was applied to the PVDF membrane, and the blot was exposed immediately to the imaging system. The image was acquired for 3-5 minutes with every 10-30 seconds interval. The band intensity is quantified and analyzed in Image J software (http://imagej.nih.gov), and the statistical significance was calculated.

#### Drug efflux assay

Drug efflux by MDR1 was measured with the Multidrug Resistance Direct Dye Efflux Assay kit (Merck Millipore; ECM 910) according to the manufacturer’s protocol. Untreated or Peptide treated cells (1×10^6^/ml) were incubated with the molecular probe DiOC2(3) (1:1000) for 15 min at 4°C in efflux buffer (20% RPMI 1640 medium, 1% BSA, 80% H_2_O). After incubation, cells were washed with the efflux buffer. An aliquot of cells was kept on ice for analysis of total dye uptake. To measure the efflux rate, an aliquot of cells was mixed with DMSO (1:1000) (diluent control) and incubated at 37°C for different time periods in efflux buffer. To check the specificity of efflux through MDR1, vinblastine (1:1000) (MDR1 specific modulator) was added to another aliquot of cells and kept at 37°C for indicated time points. Efflux was stopped by resuspending the cells in ice-cold efflux buffer, and kept on ice until analysis could be performed. Cells were then centrifuged at 200 x g for 5 minutes at 4°C and the supernatant was removed. Finally, the cells were resuspended in 0.25 ml cold efflux buffer. Cell suspensions were dispensed into the wells of a black-walled 96-well plate and measured in a fluorescence plate reader at an excitation wavelength of 485 nm and an emission wavelength of 530 nm.

## Results and Discussion

In several studies before, it has been well-established that positive coactivator 4, PC4, interacts with the wild-type p53 and assists it to regulate many of the genes targeted by p53. However, very little is known about the interaction of GOF mutant p53 proteins with PC4 and its role in the gain of new functions. We chose one of the most prevalent GOF mutants, R273H, and studied its interaction with PC4 *in vitro*.

Whole cell extracts of SW480 (R273Hp53^+/+^) or H1299 (R273Hp53^−/-^) cells, either transiently or stably expressing GOF mutant R273Hp53, were immunoprecipitated with antibodies specific for p53, PC4 or normal IgG. Immunocomplexes and inputs (5% of the whole cell extracts) were then probed with either p53 or PC4 antibodies as indicated. The results demonstrated that mutant p53 can interact with PC4 (Figure S2A, Figure S2B). In previous studies (*9*), we have shown that a small peptide (PeptideAc), triply acetylated at three lysines, derived from the extreme C-terminus of p53 disrupts the wild-type p53-PC4 interaction specifically. We have used the same peptide to determine if the R273Hp53-PC4 interaction is specific and similar to that with the wild-type p53. A significantly weaker band of PC4 in Western Blot was observed when pulled down with anti-p53 antibody in the lane marked PeptideAc (Figure 1) indicating that the peptide is capable of displacing R273Hp53 from PC4. The mutant peptide, PeptideMut, which does not bind to PC4 (*8*), is unable to significantly weaken the PC4 band, indicating its inability to disrupt the R273Hp53-PC4 complex. Similar results were obtained in SW480 cells (Figure S2C). This suggests that the protein-protein interaction observed between PC4 and R273Hp53 is specific and may be similar to that of the wild-type p53.

**Figure 1:**
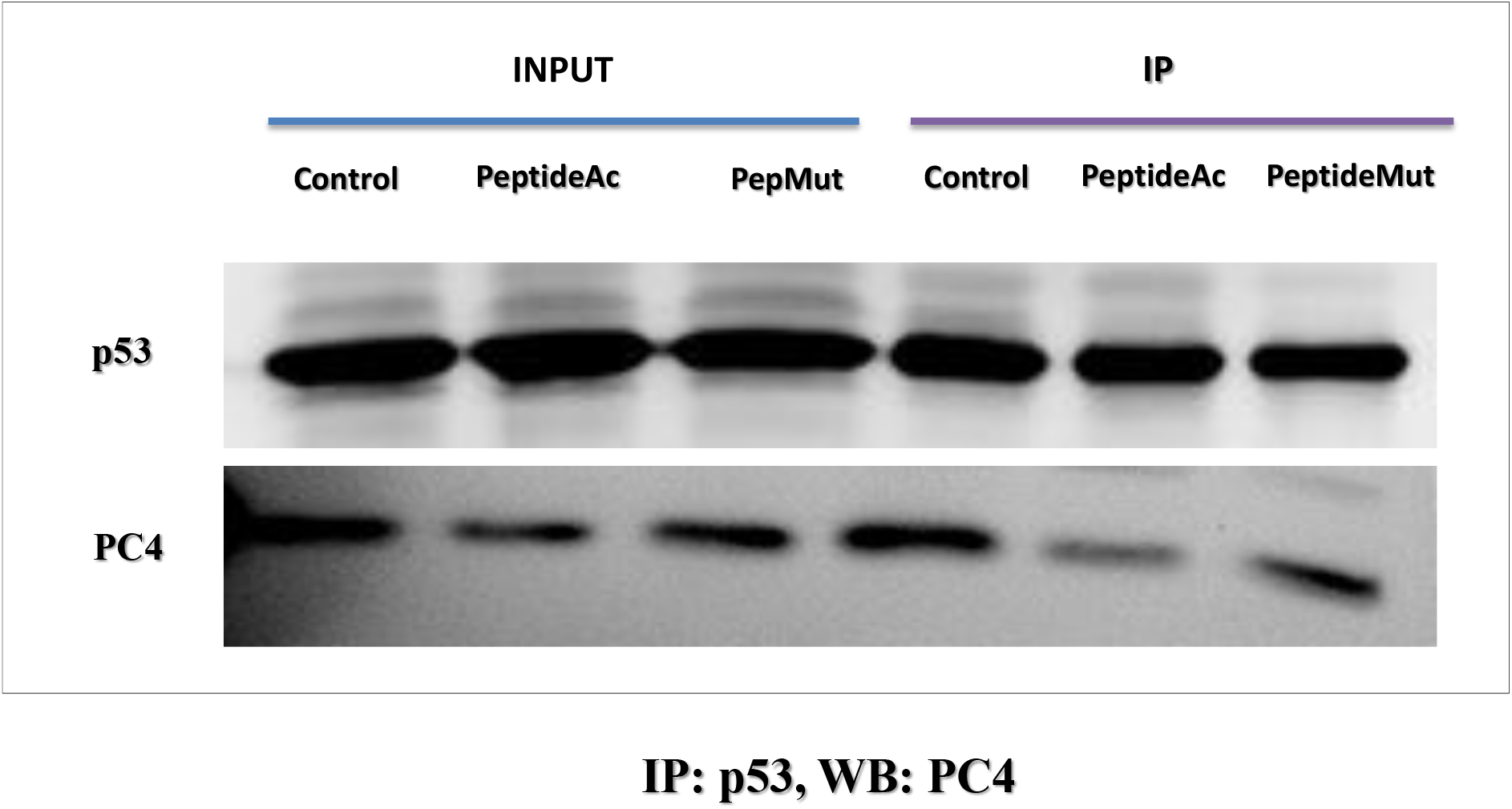
Interaction of R273Hp53 with PC4. H1299 cells bearing FLAG-R273Hp53, treated with PeptideAc or PeptideMut. Cells were treated with peptides for 12 h. Untreated cells were used as a control. Whole cell lysates were used as Input. Exogenous p53 was pulled down by the antibody stated, and western blot was performed with above-mentioned antibodies as stated. Image of 1 blot out of 3 independent experiments is given.

### R273Hp53 and PC4 are recruited to promoters of genes regulated by R273Hp53

PC4 is a chromatin-associated protein and transcription coactivator. p53 is a transcription factor whose targets are also mostly on chromatin. Thus, we have examined whether these two proteins interact on cis-regulatory elements of target genes. The presence of a highly conserved CCAAT-CCAAT-CHR module in the UBE2C promoter, which is responsible for the GOF mutant R273Hp53-mediated activation of UBE2C expression, was reported previously. Another well-known R273Hp53 target gene is MDR1. We have thus chosen regulatory regions of these target genes to explore whether both are recruited to these promoters and interact on the promoters.

We further reported that DNA damage induces the recruitment of p300, to the UBE2C promoter by GOF mutant R273Hp53 (*10*), which then promotes the acetylation of appropriate histones. p300 also acetylates wild-type p53 at position 382 and PC4 upon DNA damage (*11*). Acetylation of both p53 and PC4 enhance their interaction and their interaction with DNA, thereby up-regulating the transcriptional function of p53 (*12*). Thus, a chromatin immune-precipitation experiment was carried out in R273Hp53 harboring SW480 cells by using antibodies against p53, acetyl p53 and PC4 or pre–immune rabbit anti-sera (control IgG), in the presence and absence of a DNA damaging agent. Results showed that both p53 and PC4 could be recruited on the promoter of UBE2C. In presence of 5-FU, the recruitment of acetylated p53 got significantly enhanced, although R273Hp53 did not show a significant increase in the recruitment. PC4 recruitment to the UBE2C promoter got moderately enhanced on drug treatment (Figure 2A). Down-regulating PC4 by siRNA significantly decreased the recruitment of acetylated p53 (Figure 2B), suggesting that PC4 is crucial for recruiting acetylated R273Hp53 to the UBE2C promoter. In the reciprocal experiment, R273Hp53 was knocked down by siRNA, which significantly decreased the presence of PC4 on this promoter (Figure 2C). Thus, it appears that PC4 and R273H are recruited in a coordinated manner. A similar experiment was performed in the MDR1 gene regulatory region, which also suggests the presence of R273Hp53 and PC4 on that promoter as well (Figure 2D).

**Figure 2.**
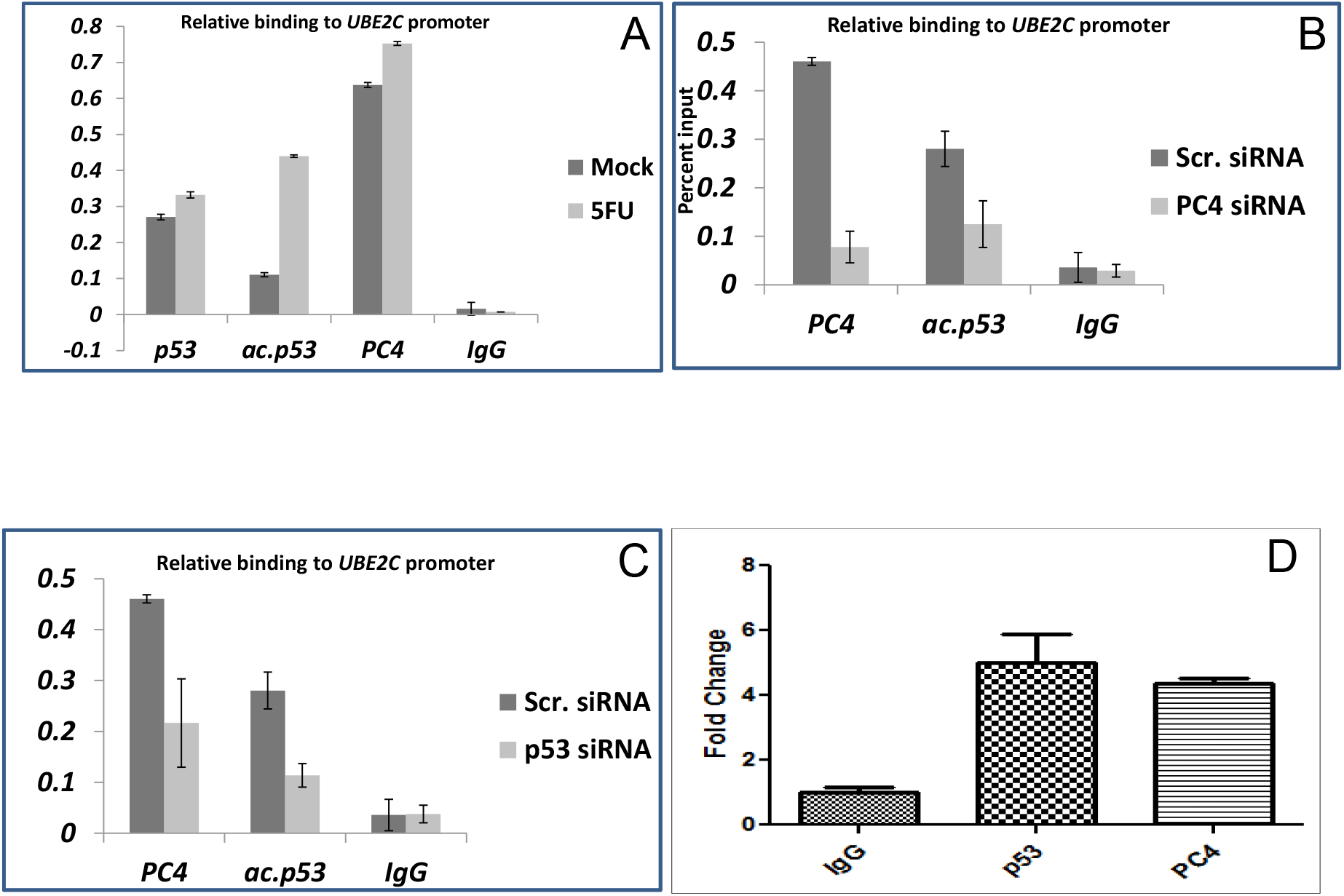
Chromatin immunoprecipitation of p53 and PC4 on two different promoters(A) ChIP assay was done in SW480 cells post treatment with or without 5-FU (10µg/ml for 16 hours) with antibodies against p53, acetylated p53, PC4 and control rabbit IgG. Precipitated chromatin was estimated by quantitative real time PCR with primers encompassing -141/+39 nt region of the UBE2C promoter relative to the transcriptional start site. Results are expressed as percent input (B) ChIP analysis was done in SW480 cells (treated with 10µg/ml of 5-FU for 16 hours), transfected with either scrambled siRNA or PC4 siRNA (80µM), with antibodies against PC4, acetylated p53 and control rabbit IgG. Precipitated chromatin was estimated by quantitative real time PCR with primers encompassing -141/+39nt region of the UBE2C promoter relative to the transcriptional start site. Results are expressed as percent input. (C) ChIP analysis was done in SW480 cells (treated with 10 µg/ml of 5-FU for 16 hours) transfected with either scrambled siRNA or p53 siRNA (80 µM) with antibodies against PC4, acetylated p53 and control rabbit IgG. Precipitated chromatin was estimated by quantitative real time PCR with primers encompassing -141/+39 nt region of the UBE2C promoter relative to the transcriptional start site. Results are expressed as percent input. (D) Chromatin immunoprecipitation at MDR1 promoter. The experiment was performed in H1299 p53 R273H cells with anti-p53 and anti-PC4 antibody. Rabbit pre-immune IgG was served as negative control. Sequences of the primer pair are shown in Table S1.

We have also attempted to determine the functional roles of recruitment of PC4 to the UBE2C promoter by knock-down and over-expression of PC4. Figure S3 shows the effect of knock-down and over-expression of PC4 on the expression of the UBE2C gene. Knock-down of PC4 in SW480 cells leads to significantly lower expression of the UBE2C gene (Figure S3A), whereas over-expression leads to modest enhancement of expression (Figure S3B). Clearly, recruitment of PC4 to UBE2C promoter leads to regulation of expression of the UBE2C gene.

In a previous study, it was shown that the GOF–mutant R273Hp53, deficient in its ability to bind to the DNA by itself, piggybacks on the transcription factor NF-Y and recruited to the CCAAT elements, thereby enhancing UBE2C expression. It was also previously reported that p300 gets recruited on the EFNB2 promoter, followed by acetylation of mutant p53 and resulting upregulation of EFNB2 expression (*13*).

We thus explored whether the peptide can disrupt R273Hp53 interaction with PC4 on promoters, which may lead to abrogation of the enhanced expression of the gain-of-function related genes. Figure 3A shows the results of CHIP experiments on the UBE2C promoter. PeptideAc was able to significantly reduce the recruitment of both PC4 and R273Hp53 on the promoter of UBE2C. The mutant peptide showed no significant effect, suggesting that the effect is specific. Figure 3B shows the result of a similar experiment on the MDR1 promoter. In this case, also, there was a significant reduction in recruitment of both R273Hp53 and PC4 to the promoter. Lesser recruitment of polII suggests a decrease in promoter activity.

**Figure 3:**
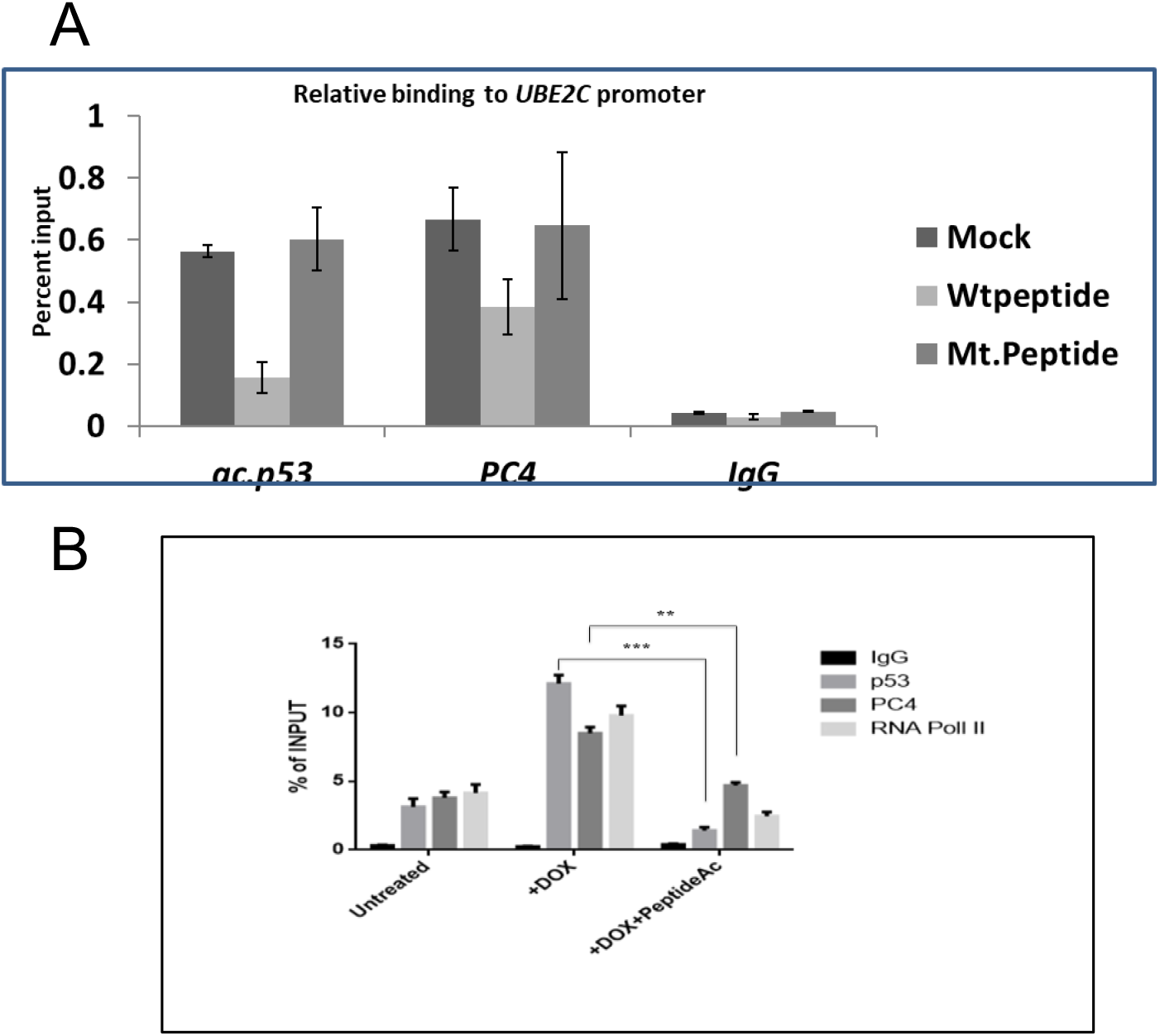
The triply acetylated wild type peptide competitively inhibits the physical interaction of PC4 and GOF mutant R273Hp53, abolishing their recruitment on UBE2C and MDR1 promoters. A) ChIP analysis was done in SW40 cells (treated with 10µg/ml of 5-FU for 16 hours) transfected in presence of either wild type or acetylated peptide with antibodies against PC4, acetylated p53 and control rabbit IgG. Precipitated chromatin was estimated by quantitative real time PCR with primers encompassing -141/+39nt region of the UBE2C promoter relative to the transcriptional start site. Results are expressed as percent input. (B) Chromatin immunoprecipitation (ChIP) was performed in H1299 cells harboring mutant R273Hp53 gene. Rabbit IgG was used as a negative control, and 5% of the total lysate was used as the input control. RNA pol II served as the positive control. Cells were treated with DOX for 12 h or PeptideAC for 12 h followed by DOX for another 12 h. Results were compared with untreated cells. The DNA-Protein complex was immunoprecipitated by using anti-p53antibody and anti-PC4 antibody. Quantitative-PCR was performed with immunoprecipitated DNA. Primers were designed from the MDR1 promoter region (Table S2). All the experiments were performed thrice, and data points are shown as mean ±SEM. Standard Error was estimated from triplicate wells. *, **, and *** indicate P≤0.01, P≤0.001, and P≤0.0001, respectively.

As it was clear from the preceding experiments, that R273Hp53 is recruited to selected promoters, perhaps by piggybacking on other transcription factors, and interacts with PC4 resulting in the stabilization of the protein complex on selected promoters. This complex is capable of upregulating at least two genes, UBE2C and MDR1. The interaction can be disrupted by the small acetylated peptide in a specific manner. We then further explored the functional effects, such as drug resistance, in the context of gain-of-function properties.

Figure 4 shows the expression of the *MDR1* gene in three different cell lines and the effect of the peptide on its expression level. We also use the substituted peptide (PeptideMut), which does not bind to PC4, as a control. The *MDR1* gene was expressed at a higher level upon treatment with Doxorubicin. Upon the peptide treatment, there was very significant reduction in the level of MDR1 mRNA in H1299(p53-/-) cells transfected with the *R273HTP53* gene. The substituted peptide had no effect on the mRNA levels. On the other hand, H1299(p53-/-) cells transfected with the wild-type TP53 gene did not show any reduction upon treatment with the peptide. The experiments clearly indicate that the mutant R273Hp53 protein forms a complex with PC4 at UBE2C and MDR1 promoters, upregulating their expression. It also shows that the loss of this interaction leads to the abrogation of this upregulation. Panc1 cells, which bear the same mutation in the chromosomal *TP53* gene, also show a modest decrease in MDR1 expression upon peptideAC treatment.

**Figure 4:**
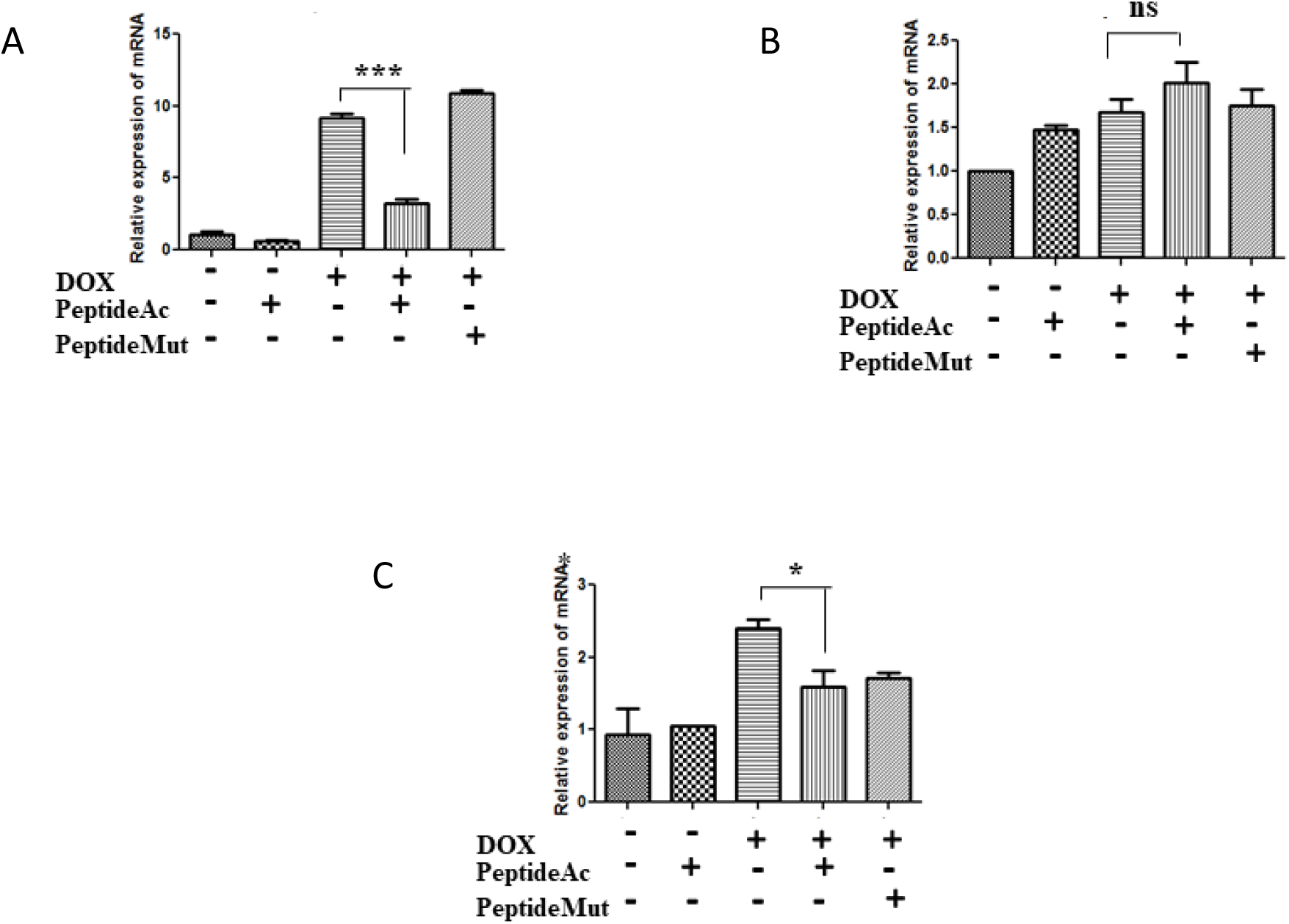
MDR1 mRNA expression of H1299 bearing (A)R273Hp53, (B) wtp53, and (C) Panc1 cells (having R273H mutation in its TP53 genes). Cells were treated with PeptideAc or PeptideMut for 24 h followed by DOX for another 24 h.

The cellular effect of the peptide was tested on H1299 cells bearing different TP53 variant genes. The MDR1 gene product is a transmembrane glycoprotein (P-glycoprotein 1) which is responsible for the efflux of many foreign substances in the cell, including many drug molecules (*14*). Over-activity of this pump often leads to pleiotropic drug resistance. Thus, we tested whether overexpression of the MDR1 gene results in higher pumping activity of the P-glycoprotein in R273Hp53 bearing H1299 cells and treatment with PeptideAC will reduce the activity of P-glycoprotein. The assay was conducted using a dye which is a substrate of the P-glycoprotein (DiOC2(3)) (*15*). Figure 5 shows that the efflux of the dye from the cell was largely inhibited when the cells were treated with PeptideAc. When another drug, vinblastine, a substrate of the P-glycoprotein(*16*), was added along with DiOC2(3), a significant reduction of the efflux of the dye was observed, suggesting that the reduced efflux seen upon PeptideAC treatment is a result of reduced pumping activity of the P-glycoprotein. When the cells were incubated at 4°C, enhanced retention of the dye was also observed. At this temperature activity of the membrane pumps will be minimal and the enhanced retention is consistent with efflux being a result of the activity of the membrane pumps. Since PeptideAc reduces efflux, it is anticipated that drugs, such as doxorubicin, may have an enhanced killing effect on R273Hp53 bearing cells, treated with PeptideAc. Right panels of Figure 5 show the viability of cells treated with PeptideAc and doxorubicin. In H1299 cells bearing the R273HTPp53 gene, a significant reduction of cell viability was observed in comparison with cells treated with the mutant peptide, PeptideMut. However, in H1299 cells, either bearing the wild-type allele of TP53, or having no functional TP53 gene, no reduction in cell viability was observed for either treatment. These observations indicate the ability of PeptideAc to sensitize cells to doxorubicin killing in R273HTP53 bearing cell lines.

**Figure 5.**
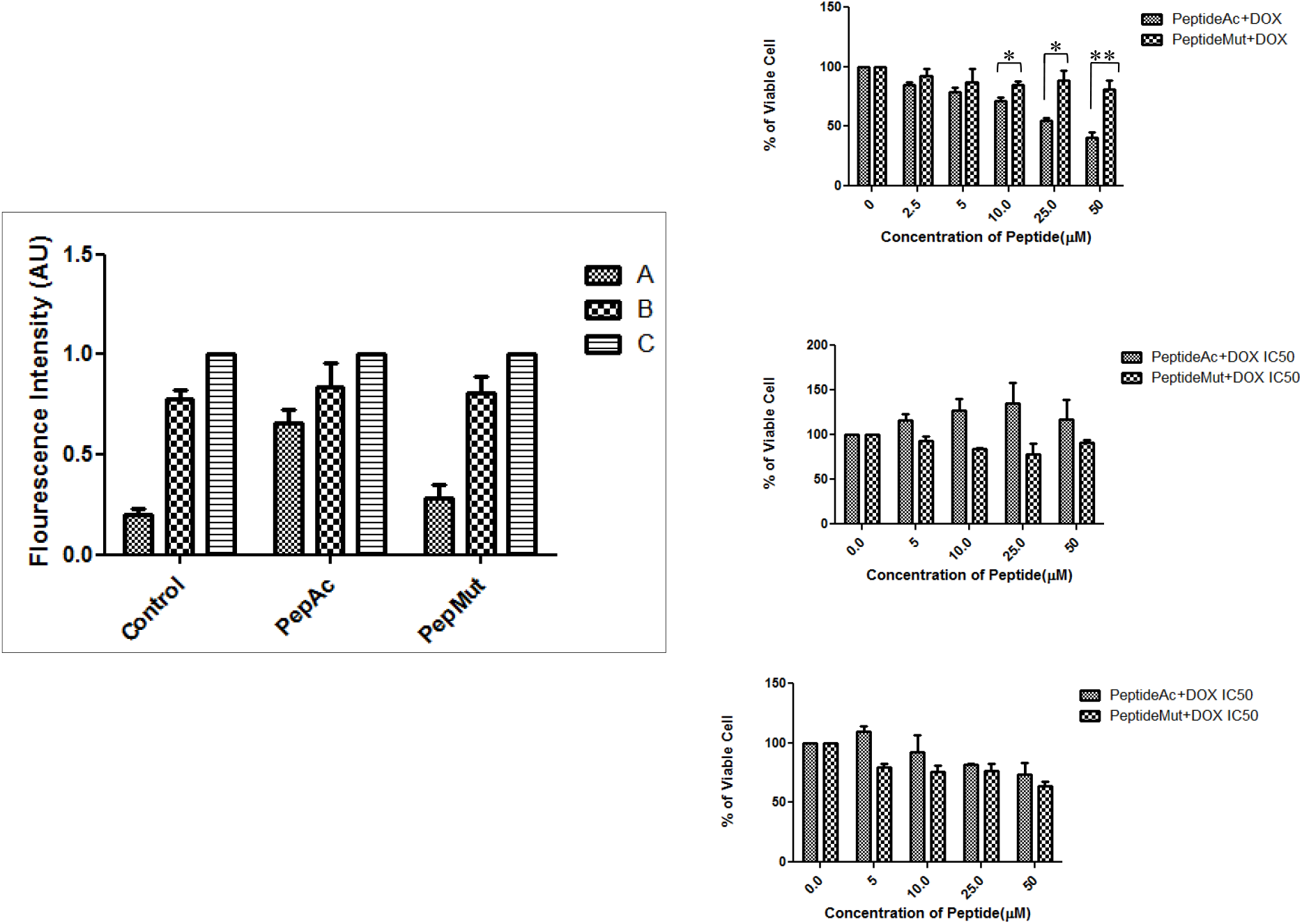
Left panel, drug accumulation and efflux assay. The accumulation of DiOC2(3) (a substrate of MDR1) was measured in untreated, R6-NLS-p53(380–386-3Ac) (PeptideAc) or R6-NLS-p53(380–386-m) (PeptideMut) treated cells by fluorescence plate reader. The retention of DiOC2(3) after a 3 h drug-free efflux was measured in three different conditions. (A) Incubation at 37°C (B) incubation with another MDR1 substrate Vinblastin at 37°C and (C) incubation at 4°C (control). Right panel,Cell viability upon DOX treatment on cells treated with R6-NLS-p53(380– 386-3Ac) (PeptideAc) or R6-NLS-p53(380–386-m) (PeptideMut) for 24 h followed by DOX (at IC_50_) for 72 h. Cells treated with DOX (IC_50_) alone have been used as control. (E) H1299 cells transfected with R273Hp53 gene, (F) H1299 cells transfected with WTp53 gene, and (G) H1299 cells (p53 null).

### The effect of PeptideAc is mutant specific

Previous experiments described in this article suggest that PeptideAc acts on R273H gain-of-function mutant p53 with high specificity. It is well-known that different gain-of-function mutants of p53 often have different effects on gene expression (*17, 18*). It is possible that different mutants may associate with different factors giving rise to different patterns of gene expression(*4*).

Thus, we explored whether another gain-of-function p53 mutant behaves similarly to R273Hp53. We chose the R248W mutant p53, which is abundantly observed in many tumors. Figure 6 shows the expression of the MDR1 gene in the presence and absence of PeptideAc in doxorubicin treated cells (H1299 cells stably transformed with R248WTPp53 gene). As observed with the R273H mutant, treatment with doxorubicin leads to about 2.5-fold enhancement of MDR1 mRNA concentration. However, in contrast to cells bearing the R273HTPp53 gene, treatment with PeptideAc had a very modest effect on the MDR1 mRNA levels. A similar effect was also observed with the PeptideMut, indicating the effect is not likely to be specific. We have also tested the viability of the cells upon treatment with doxorubicin in the presence of PeptideAc or PeptideMut. No significant difference was observed between these treatments and both were similar to the control (Figure 6). Thus, it appears that PC4 may not be a general cofactor for all GOF mutants, leading to mutant-specific effects.

**Figure 6.**
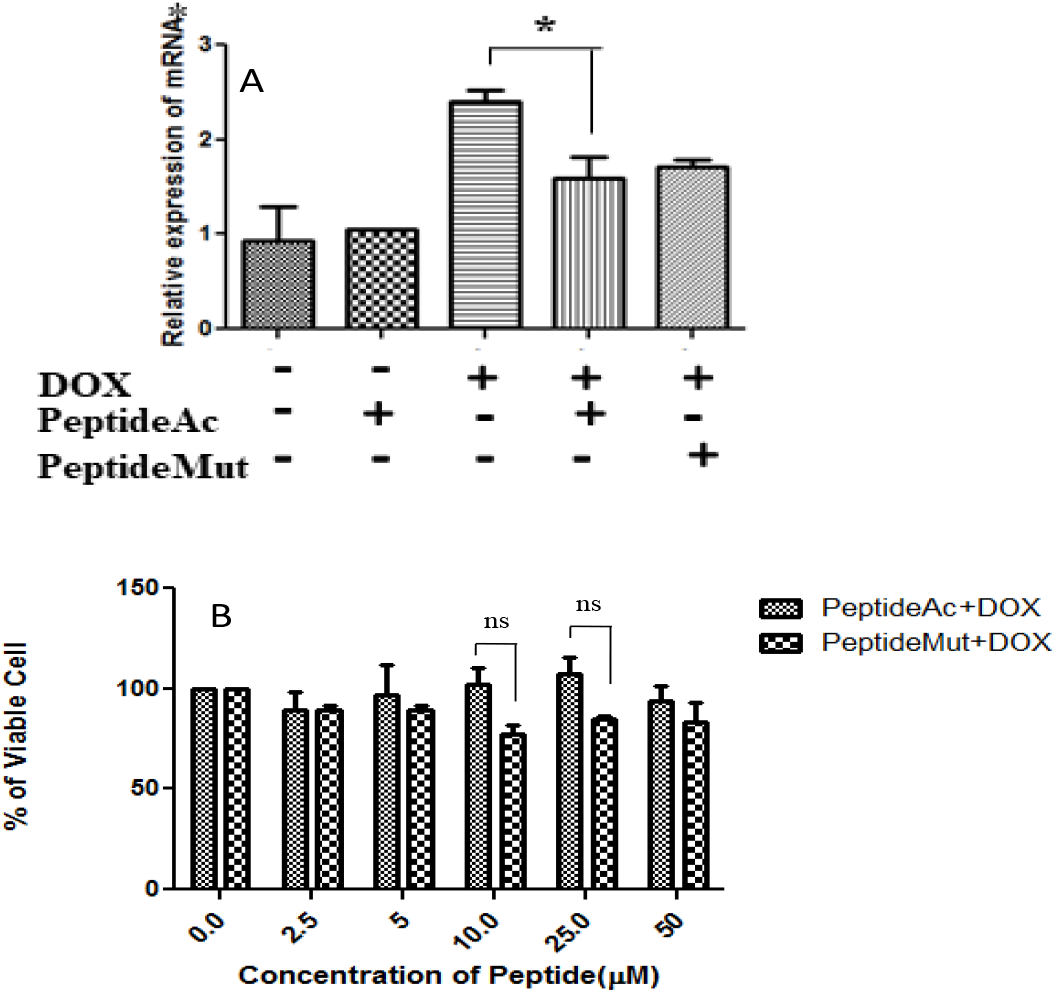
Effect of PeptideAc on H1299(-/-) cells bearing R248WTP53 gene. (A) The expression of MDR1 gene on upon treatment of PeptideAc and PeptideMut. Cells were treated with R6-NLS-p53 (380–386-3Ac) (PeptideAc) or R6-NLS-p53 (380–386-m) (PeptideMut) for 24 h followed by DOX for another 24 h. (B) The viability of the H1299(-/-) cells bearing R248WTP53 gene upon doxorubicin treatment previously treated with PeptideAc and PeptideMut.

Tumors containing GOF mutant p53 proteins pose a serious challenge to offering viable therapeutic options. A major characteristic of these tumors is the pleiotropic resistance to drugs, making chemotherapies difficult. Reversal or abrogation of the drug resistance is a desired goal for the treatment of this class of tumors. The R273H mutation in the TP53 gene occurs in many tumors with significant frequency. Many attempts have been made to target GOF-mp53 proteins (*19*). We have identified PC4 as a novel target for R273Hp53. PC4 interaction with p53 has been shown in this study to be crucial for the upregulation of the MDR1 gene, which is often the root cause of pleiotropic drug resistance. Disruption of this interaction with a peptide reverses the drug resistance, making the cells sensitive to chemotherapeutic agents. This opens up the possibility of targeting PC4 in tumors bearing the R273H mutation. In the future, the peptide may be replaced by small molecules having similar functions.

## Supplementary Information

**Table S1:**
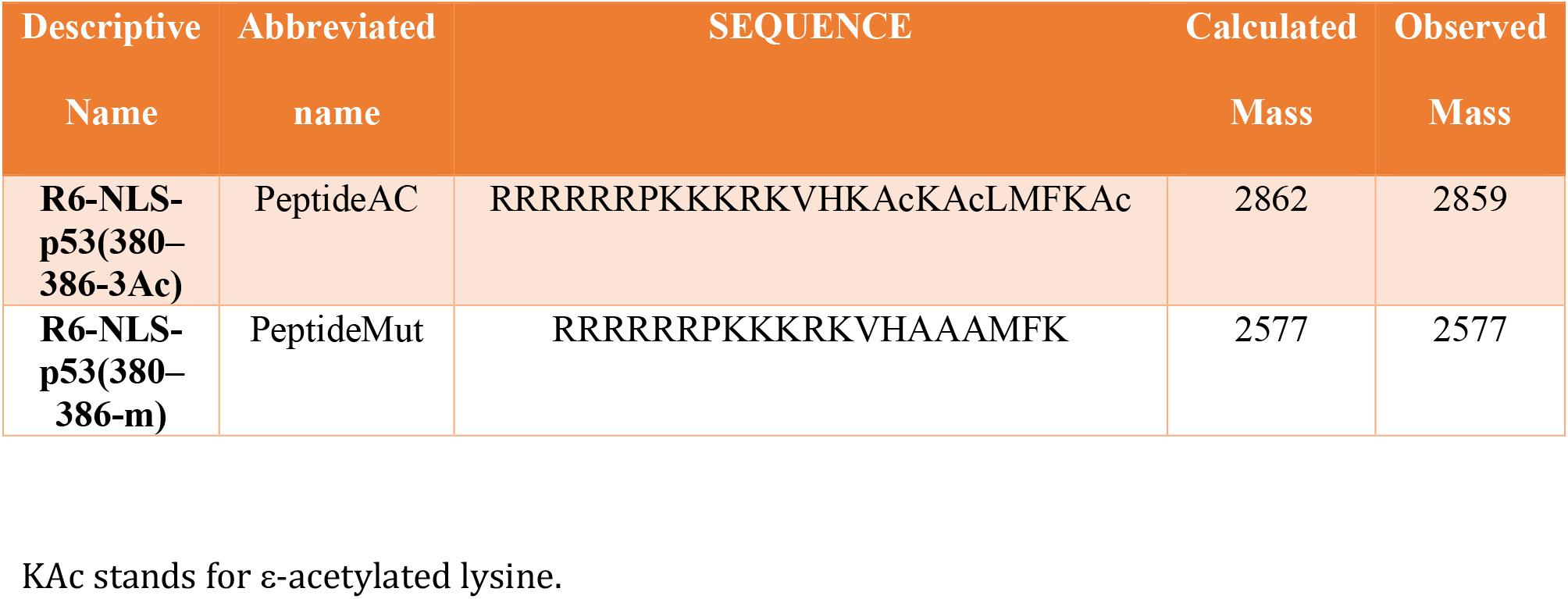
Characterization of peptides used

**Table S2.**
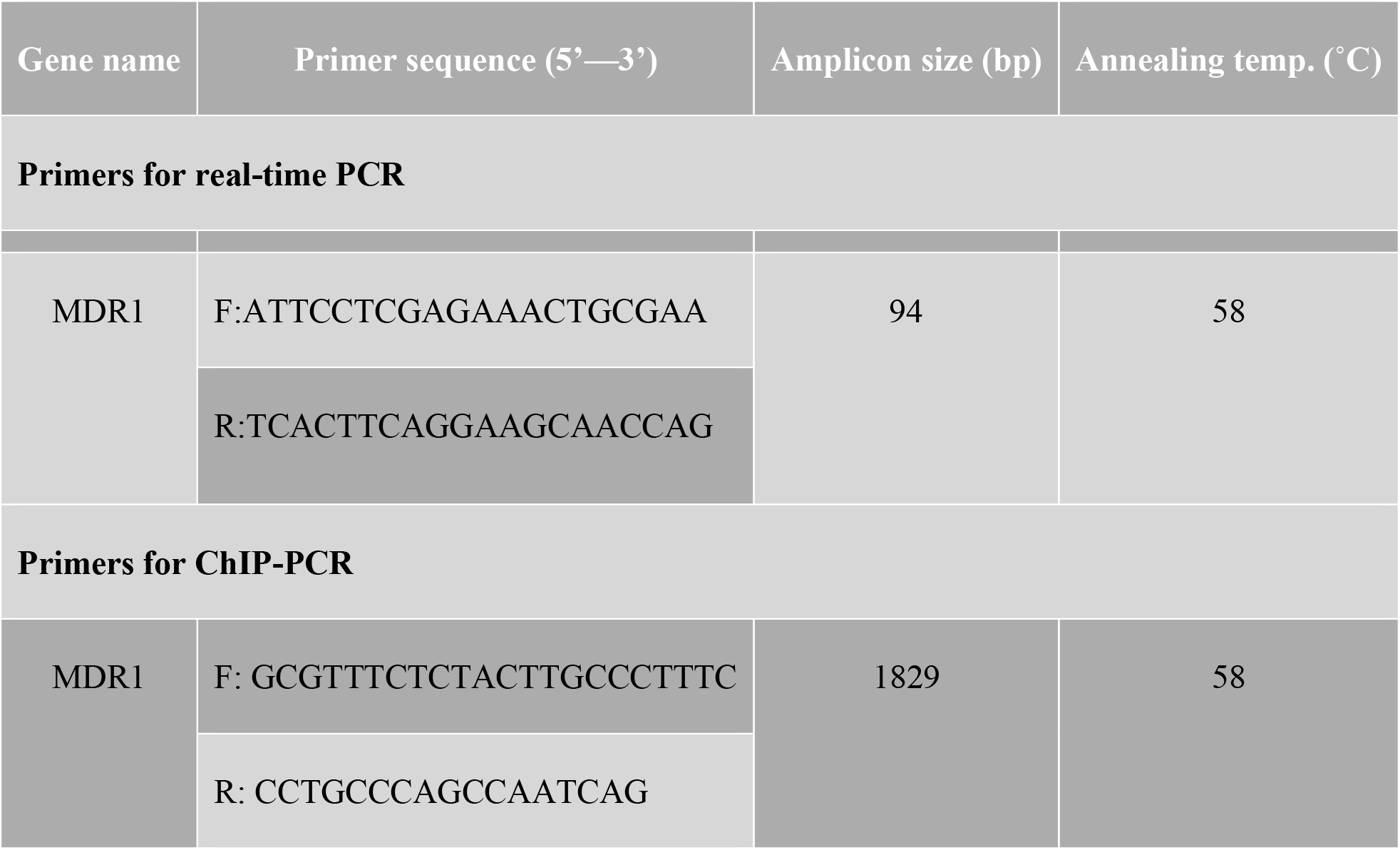

**Figure S1.**
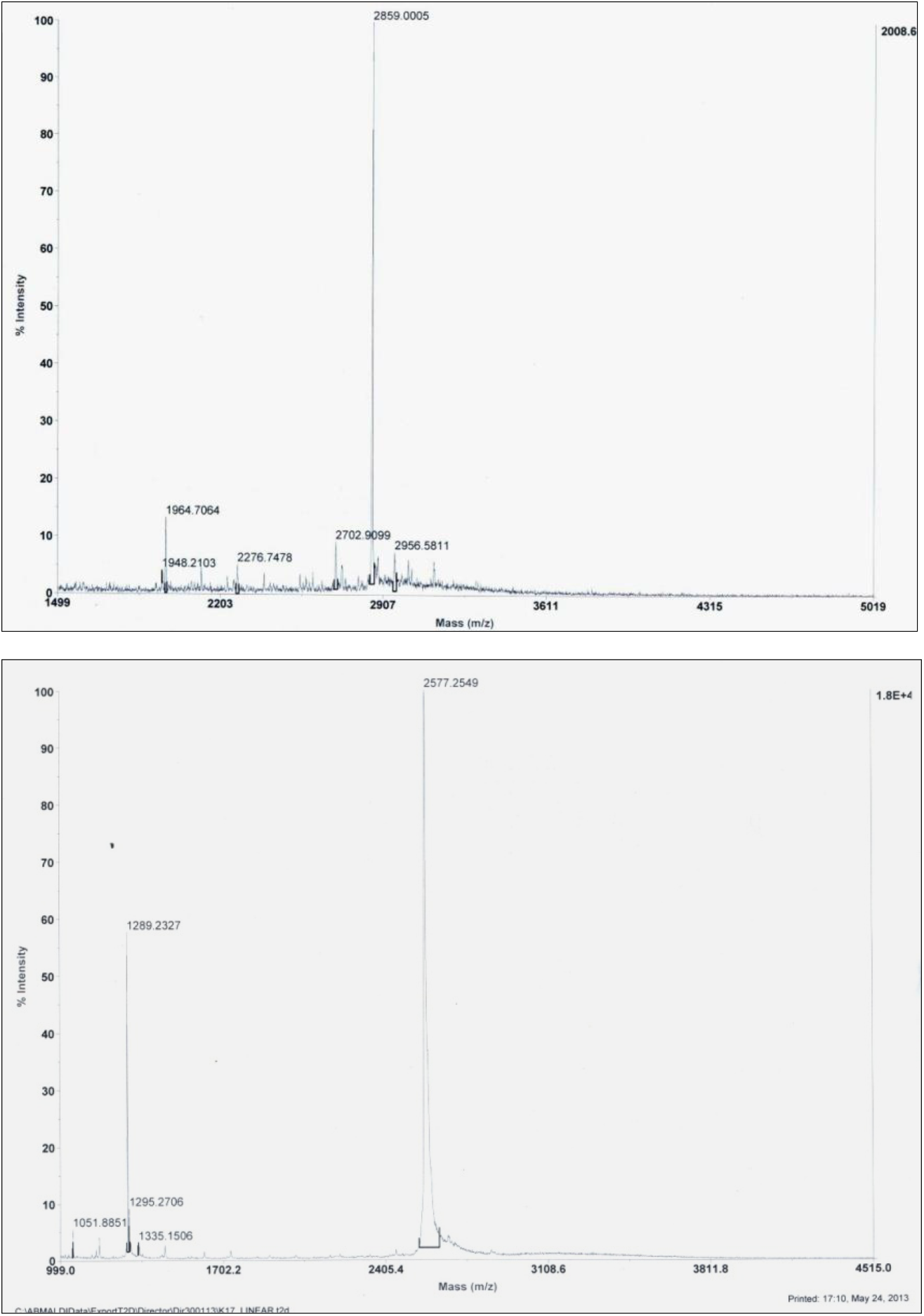
Top panel, Maldi-TOF mass spectra of PeptideAC. Bottom panel, Maldi-TOF mass spectra of PeptideMut.

**Figure S2:**
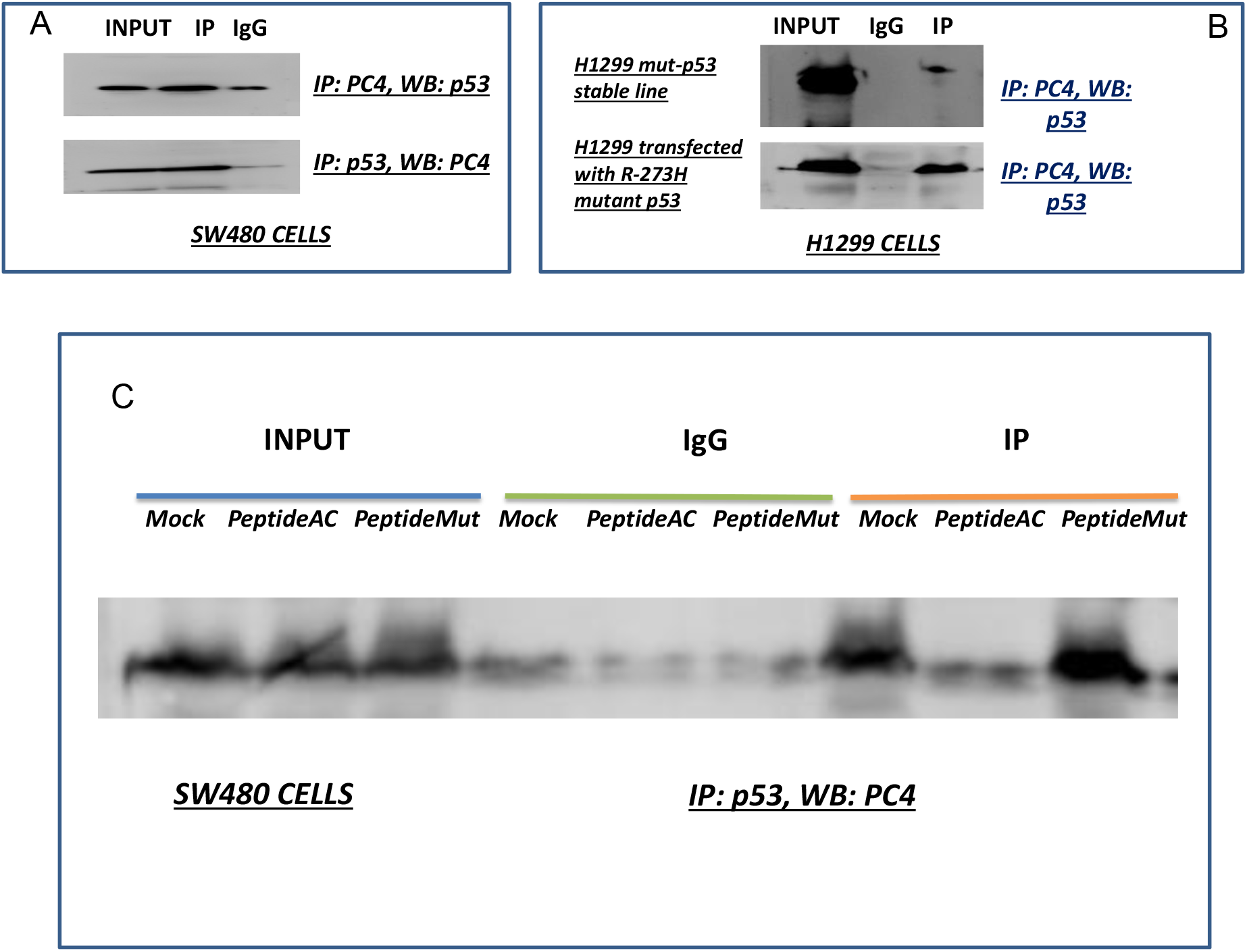
Interaction of R273Hp53 with PC4 in different cells. (A) Immunoprecipitation of R273Hp53-PC4 complex from SW480 cells. The cell lysates were either precipitated with anti-p53 antibody and probed with anti-PC4 antibody or the other way around. (B) Immunoprecipitation of R273Hp53-PC4 complex from H1299 (p53^−/-^) cells, which contained either stably transfected R273Hp53 or transfected with plasmids containing R273Hp53. The cell lysates were precipitated with anti-PC4 antibody and probed with anti-p53 antibody. (C) PeptideAc disrupts the mutant p53 and PC4 interaction ex-vivo. Whole cell extracts of SW480 cells treated with either wild type (wtp) or mutant peptide (mtp) were immunoprecipitated (IP) with antibodies against p53 or normal IgG. Immunocomplexes and input (10% of the whole cell extracts) were probed with antibodies against PC4 by Western blot analysis. The experiments were performed once.

**Figure S3:**
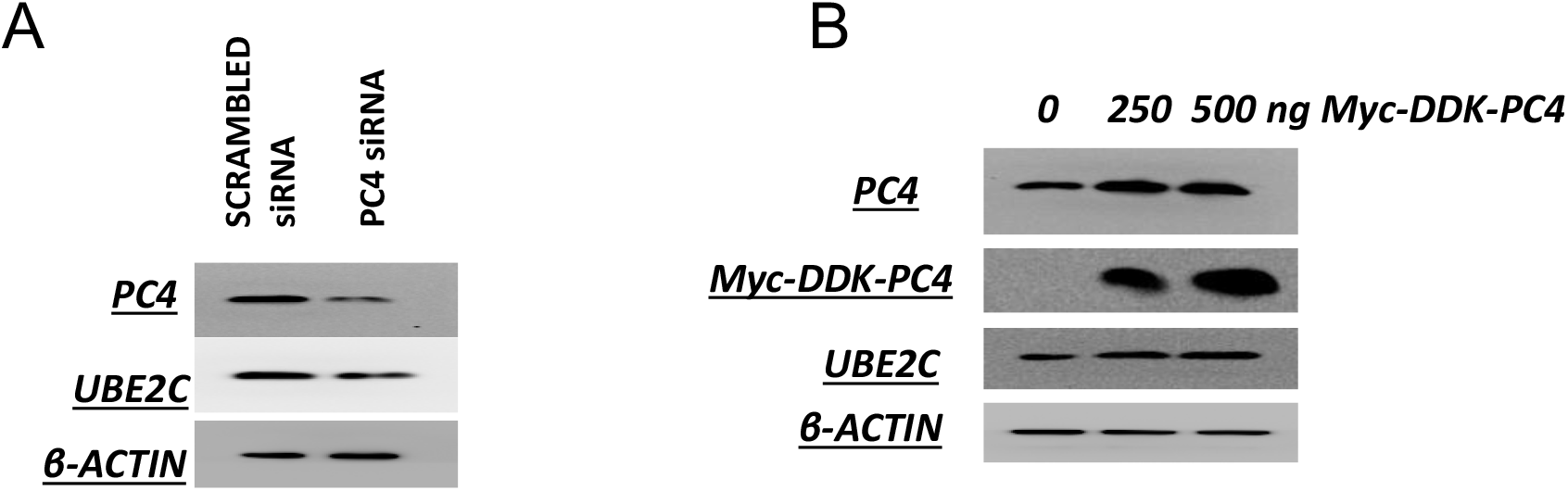
Effect of PC4 levels on UBE2C expression(A) SW480 cells were transfected with either SCRsiRNA or PC4 siRNA and western blot analysis was done with antibodies against PC4, UBE2C and β-actin. (B) SW480 cells were transfected with increasing concentrations of myc-DDK-PC4 expression plasmid and western blot analysis was done with antibodies against PC4, DDK, UBE2C and β-actin.

